# Conjoint specification of action by neocortex and striatum

**DOI:** 10.1101/2023.10.04.560957

**Authors:** Junchol Park, Peter Polidoro, Catia Fortunato, Jon Arnold, Brett Mensh, Juan A. Gallego, Joshua T. Dudman

**Affiliations:** Janelia Research Campus, Howard Hughes Medical Institute, Ashburn, VA 20147; Department of Bioengineering, Imperial College London, London W12 0BZ

**Keywords:** Neuropixels, Striatum, Motor Cortex, Motor control, Reinforcement Learning, Action specification, Action selection

## Abstract

The interplay between two major forebrain structures - cortex and subcortical striatum - is critical for flexible, goal-directed action. Traditionally, it has been proposed that striatum is critical for selecting what type of action is initiated while the primary motor cortex is involved in the online control of movement execution. Recent data indicates that striatum may also be critical for specifying movement execution. These alternatives have been difficult to reconcile because when comparing very distinct actions, as in the vast majority of work to date, they make essentially indistinguishable predictions. Here, we develop quantitative models to reveal a somewhat paradoxical insight: only comparing neural activity during similar actions makes strongly distinguishing predictions. We thus developed a novel reach-to-pull task in which mice reliably selected between two similar, but distinct reach targets and pull forces. Simultaneous cortical and subcortical recordings were uniquely consistent with a model in which cortex and striatum jointly specify flexible parameters of action during movement execution.

**One sentence summary:** Motor cortex and subcortical striatum act in concert to specify the movement parameters of a reach-to-pull action in mice.

## Introductions

Mammals have multi-jointed limbs with many degrees of freedom enabling highly flexible, complex, and dexterous actions. It has long been appreciated that these many degrees of freedom pose daunting control challenges for the nervous system and any given action is characterized by the many degenerate solutions that can all achieve a given motor goal (*1*). For example, for an animal to successfully reach out to grasp a piece of food the forelimb can take many different trajectories each implemented by a complex pattern of activation across many muscles. While our understanding of the processes underlying the planning and execution of actions is understandably incomplete, a number of principles and a relatively detailed outline has emerged (*2–5*). The canonical functional sequence requires a target to be selected, converted via a sensorimotor coordinate transform into egocentric coordinates, an effector and kinematic trajectory planned, followed by online control of movement execution. In a reach-to-grasp action this sequence might involve identifying the location of a piece of food, computing a vector from the current hand location to the food, choosing to reach from below to avoid an obstacle, and reaching slowly with high co-contraction to accurately grasp the food. One principled distinction in these functional processes is between ‘selection’ operations that reduce a set of *discrete* possibilities to a single option and ‘specification’ operations that set *continuous* parameters governing execution (*2*, *4*, *6*). For example, one may select which piece of food to target or which forelimb to use, in distinction from specifying a particular speed of movement or tightness of grasp.

Seminal work has mapped this abstract motor planning and execution sequence onto a distributed and partially dissociable set of brain areas thought to implement these functional processes. For example, in neocortex the selection of a target and transformation into egocentric coordinates during motor planning maps well onto posterior parietal cortical areas (*3*). Action selection maps onto processes in premotor cortical areas of the medial wall of frontal cortex such as anterior cingulate areas in the primate (*2*, *4*, *5*, *7*) and likely onto homologous areas of cingulate and secondary motor cortex in rodents (*8*). Action specification, in distinction, maps onto the primary motor cortex (MOp) (*9–12*). One particularly strong piece of evidence for specification in MOp is the relatively enhanced encoding of movement trajectories (kinematics) and forces (kinetics) in MOp relative to reduced decoding of movement details in cingulate areas (*7*). Moreover, disruption of MOp function can produce profound deficits in voluntary forelimb control (*10*). However, the cortex does not act alone, but rather interacts with subcortical structures that are critical for flexible, dexterous actions (*6*, *10*, *13–15*).

The striatum (STR) is a subcortical forebrain nucleus (*16*) that receives convergent input from MOp, secondary, parietal and frontal cortical areas and projects to downstream nuclei of the basal ganglia (*16–18*). The dorsal striatum (dSTR) is known to be critical for the establishment of motor skills guided by reinforcement teaching signals conveyed via dense innervation from midbrain dopamine neurons (*6*, *13–16*, *19–22*). In analogy to the discrete action spaces of canonical reinforcement learning (RL) models (*23*), dSTR has long been proposed to play a specific and circumscribed role in action selection (*19*). RL models used in neuroscience (*e.g.* (*24–26*)) tend to reduce the control of action to a selection process at the moment of movement initiation or at each moment of an unfolding limb movement (*27*). As a consequence, much focus in experimental work on basal ganglia has been on analyzing the moment an experimenter observes an animal initiate one of a discrete set of actions (*e.g.* move left or right; (*24*, *25*, *28*, *29*)). These data have suggested that dSTR may mediate premotor-like (*e.g.* as seen in cingulate areas) action selection processes. However, work that has begun to quantify continuous temporal and spatial variability of movement execution has increasingly implicated dSTR in action specification processes functionally equivalent to functions of primary motor cortex (*6*, *30–33*).

Two specific limitations in prior work have made it difficult to resolve the function of dSTR in action selection and execution. First, there has been little work recording dSTR simultaneously with premotor and primary motor areas during execution of flexible and dexterous movements. Thus, it has been difficult to assess whether dSTR activity is more consistent with activity predicted for action selection functions of medial wall premotor areas or action specification functions of primary motor areas. Such comparisons arising from whole brain functional imaging or paired electrophysiological recordings in the primate neocortex have been critical to such distinctions (*2*, *7*). Second, while electrophysiological measurements spanning cortical and subcortical structures are now tractable in the rodent, prior behavioral paradigms in rodents have focused either on a single stereotyped action (*e.g.* (*32*)) or continuously varying actions (*e.g.* (*34*, *35*)) without a requirement for both discrete selection of movements and continuous specification of movement execution parameters. Thus, in rodents the critical behavioral contrasts have not been combined with multiarea recordings. In the current study we address both of these limitations.

### Distinguishing predictions

In developing the behavioral paradigm used here let us first clarify a key point: in the case of highly distinct actions, selection and specification hypotheses make essentially indistinguishable predictions about neural activity (*6*). Take for example a comparison of activity between orienting left or right or initiating a rearing movement vs locomotion (*36*, *37*). According to a selection hypothesis these actions are associated with distinct active ensembles of neurons exclusively in dSTR (*28*) although under other variants distinct ensembles of active neurons are found throughout the motor system (*38*). Specification operations make the same prediction. Different sets of muscles are being controlled with distinct spatiotemporal dynamics thus requiring distinct active neural ensembles (*6*, *31*). This could lead one to suggest the differences between these models are therefore merely “semantic.” However, comparing two very distinct actions, although pervasive in previous studies, is a specific case in which selection and specification models are indistinguishable, but there are different experimental comparisons where the models can be readily distinguished. Perhaps paradoxically, comparing neural activity across actions differing only modestly in their kinematic or kinetic parameters makes much better distinguishing predictions (Fig. 1; Supplemental Fig. 1).

**Figure 1.**
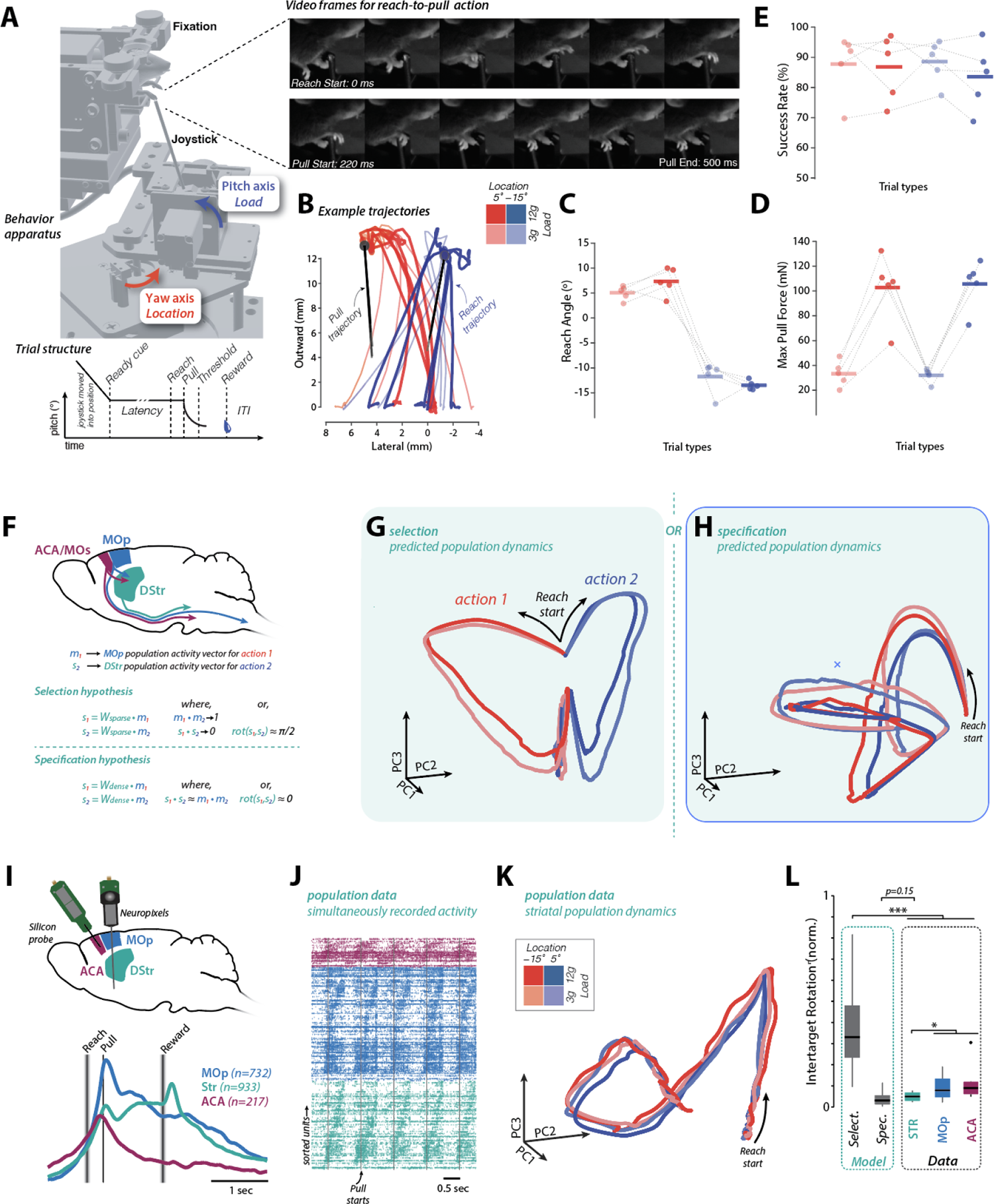
Reach-to-pull task and analysis design to distinguish predictions of specification and selection functional accounts. **A** 3D rendering of robotic joystick apparatus indicating the key degrees of freedom and motorized control of pitch and yaw axes. Right, images from dual high speed camera video of performance. Lower, timeline for individual trials. Trials were composed of 4 trials types that varied between low and high pull force (color intensity) and reach target location (blue vs red). **B** Example trajectories of successful reaching movements for each target (red and blue lines) and pull direction (black lines). **C** Blockwise average observed reach angle as a function of joystick target position. **D** Blockwise average observed pull force as a function of joystick load condition. **E** Performance is comparable across all 4 pull force + reach target conditions. **F** Schematic of key brain areas studied. Lower, quantitative description of a key difference between predictions of selection and specification hypotheses. See text for details. **G-H** Analysis of simulated encoding models developed to provide quantitative predictions from a selection model **(G)** or specification model (H) formulation. Activity was projected onto the first three principal components (PCs). **I** Schematic of recording configurations used. Lower, mean perievent time histograms (PETHs) of mean activity aligned to the initial of the in each major area targeted. **J** Raster plots of 5 sequential, concatenated trials for simultaneously recorded single units. Units are sorted according to peak time of activity modulation of PETH within trial. **K** Dynamics of population activity of striatal activity for all sessions projected onto the first three PCs of activity as in **G-H.** L Population dynamics across trial conditions were compared by quantifying the amount of rotation required to transform the population activity trajectory from one target location onto the other for all models and recordings from individual brain areas. Statistical significance was evaluated using multiple comparison corrections. *: p<0.05, **: p<0.01; ***: p<0.001.

In order to see why this is the case, consider a common task - reaching to one of two possible targets that are far from each other - and two paradigmatic models: (1) a selection model in which discrete actions (reach to target A vs B) are reflected in non-overlapping neural ensembles and (2) a specification model with continuous parameterization of action in the neural ensemble activity (Supplemental Fig. 1). To visualize differences in these distinct model formulations it is useful to visualize the projection of neuronal population activity onto dimensions that capture the majority of variance (principal components; PCs). As noted above, for very distinct actions (reaching to targets separated by 180°) one observes well separated population activity trajectories under either model (Supplemental Fig. 1). Contrast this with the trajectories expected for two similar actions differing by only, say ∼20°. A selection model is powerful because it predicts distinct neural ensembles corresponding to each action regardless of how distinct the kinematics are - thereby allowing reliable mapping of the selected action onto largely independent neural ensembles (*i.e.* see Figure 5 in (*28*)^1^). Under a specification model, however, the difference in ensemble activity between similar actions is small. When neuron ensembles are tuned to the range for a continuous parameter of an action (we consider canonical cosine tuning to direction (*40*), but the exact encoding details are not critical) a small change in movement direction accounts for little variance in population activity (Supplementary Fig. 1). As a result, the leading PCs will now be dominated by the temporal dynamics of activity common across conditions (*41*, *42*) and the trajectories of the leading PCs will exhibit small differences^2^ in distinction from the predictions of a selection encoding model (Fig. 1G). Thus, a behavioral paradigm in which an animal is reliably selecting between actions with small parameter variation can be critical to disambiguate specification from selection.

### Behavioral paradigm and neural recordings

To achieve this goal, we engineered a new joystick apparatus for head-fixed mice (Fig. 1A) (*43*, *44*). The joystick was robotically positioned at one of two equidistant yaw angles (+5°/–15°; ∼20 mm radially from hand rest) and independently commanded to force setpoints of 3g and 12g (see Methods). The mouse was required to reach out to the correct yaw target location, grab the joystick, and pull with sufficient force along the pitch axis to displace the joystick past a distance threshold (∼1 cm). Trials were divided into blocks dissociating joystick yaw location and/or force requirements along the pitch. In each session, mice completed 8 blocks, 2 repetitions of each of the 4 conditions (Fig. 1A-E; location +5°/–15°, force set point 3g/12g, Supplementary Video. 1).

Analysis of reach-to-pull behavior revealed a double dissociation in reach trajectory angle across joystick location and applied pull force (repeated-measures ANOVA; reach angle, *F*_3,12_=135.7, *P*=8.25×10^-8^; pull force, *F*_3,12_=57.93, *P*=3.30×10^-6^; Fig. 1B-E) while maintaining levels of performance (>80% of trials successfully completed) comparable to reach-to-grasp tasks with a single target location (*44–47*). Importantly, after training, performance did not differ systematically across trial types nor between individual mice (repeated-measures ANOVA, *F*_3,12_=1.33, *P*=0.32).

### Inactivation of either MOp→STR or MOs→STR projections impairs reach-to-pull performance

As noted above, the dorsal striatum receives input from both primary motor cortex and frontal premotor areas (*17*, *18*, *48*, *49*). To confirm that both sources of corticostriatal input are critical for performance of this task, we examined how silencing primary motor (MOp→dSTR) or secondary motor/anterior cingulate (MOs/ACA→dSTR) projection neurons affected task performance. For optical silencing of projection neuron populations, we used intrastriatal injection of rAAV2-retro into a dorsal striatal area known to retrogradely label both MOp and ACA in mice (*17*, *50*) (see Methods). We express a potent anion-conducting channelrhodopsin (GtACR2), previously confirmed to provide robust inactivation of cortical projection neurons (*44*), in corticostriatal projection neurons (*51*). In randomly selected trials (P=0.25) the 473-nm bilateral silencing laser was triggered during the inter-trial interval and terminated either by a trial completion or after 4 seconds. To control for visual perception of laser activation, stimuli were masked with ambient blue light (*47*). We compared the probabilities of successful reach start within 1 second after the joystick positioning (trial start). Suppression of activity in corticostriatal projections from both MOp and MOs/ACA led to a significant decrease in the probability of successful reach start (Supplementary Fig. 2; paired t test; MOp: *t*_1_=3.43, *P*=0.042; MOs: *t*_1_=5.48, *P*=9.25×10^-4^). This is consistent with a key role for ACA in movement initiation (*7*).

### Neocortical and striatal activity exhibit complex temporal dynamics during movement execution

To measure neural activity in the forebrain we acutely implanted a 384-channel Neuropixel probe in the forelimb motor cortex and the underlying dorsal through ventral striatum (Fig. 1I) (*44*). In addition, in a number of experiments we also implanted a 64-channel silicon probe in the same region of MOs/ACA that was important for movement initiation and provides dense input to dSTR in areas also innervated by primary motor cortical areas (*17*, *18*, *49*, *52*). From 5 mice we recorded 732 (avg: 81.3, sd: 44.1) forelimb MOp, 933 (avg: 103.7, sd: 23.1; dorsal: avg: 69.9, sd: 24.5) STR units over 9 recording sessions. In addition, 217 (avg: 31.0, sd: 11.7) secondary motor and cingulate (MOs/ACA) units were recorded from 4 mice over 7 sessions (Fig. 1I-J).

Both cortical and subcortical areas exhibited strong modulation throughout the reach-to-pull movement sequence and around the time of reward delivery (Fig. 1I-J). An example set of 5 sequential trials with raw spike rasters from ∼350 units recorded simultaneously from MOs/ACA, MOp and dSTR revealed highly reliable activity across repetitions of the reach-to-pull action (Fig. 1J). It is clear even from this raw data across multiple trials that there are highly structured temporal dynamics of activity in both cortex and subcortical dSTR consistent with previous descriptions in other forelimb operant tasks (*6*, *32*, *34*, *53*). We next examined the projection of striatal activity onto the leading PCs across the population of all recorded units (n=933) and separated according to the 4 trials conditions (Fig. 1K). We observed largely aligned activity trajectories that were dominated by the temporal dynamics of activity during reach-to-pull movements that did not differ from the predictions of the specification model (p∼0.15, *ranksum test*), but were outside the distribution of predictions from the selection model (p≪0.0001, *ranksum test*; Fig. 1L). Similarly, activity in MOp and MOs/ACA exhibited slightly *greater* rotations of population activity across movements to the two targets as compared to dSTR (p<0.05, *ranksum* test) even though we know that MOp in particular should not be (and is not) consistent with a selection model (Fig. 1L).

### Cortex and subcortex jointly specify movement parameters

Despite the fact that mice reliably select between closely related actions differing in their direction and force parameters, we found little evidence for highly distinct neural ensembles in dSTR across conditions. Rather, the neural ensembles were substantially overlapping and dominated by the complex temporal dynamics of activity during action execution (Fig. 1I-K). However, there are dimensions along which population activity does differ according to reach-to-pull parameters. In the case of the paradigmatic specification model, these differences are not the primary source of variance, but can be readily revealed by finding contrastive principal components (‘cPC’, see methods for details (*54*); Fig. 2A and Supplementary Fig. 3). We first examined simulated activity from a specification model to confirm that activity projected onto cPCs reveals trial-type specific geometry of activity that is not apparent in standard PCA (Fig. 2A, right). Consistent with predictions from the model, projecting population activity on cPCs revealed a distinct geometry of activity across reach-to-pull actions across target locations (Fig. 2A).

**Figure 2.**
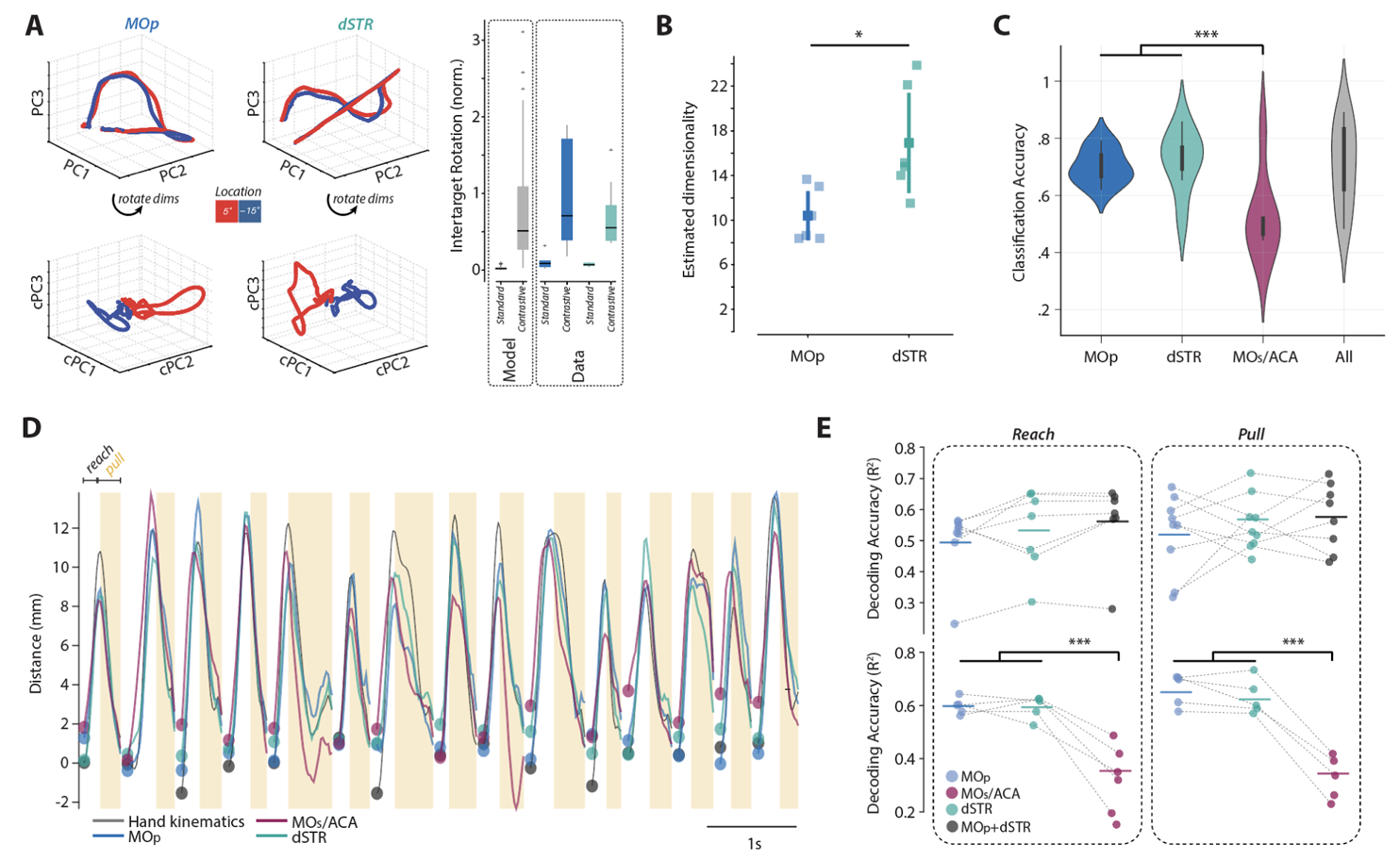
Quantifying differences in the population encoding across trial conditions. **A** Contrastive principal component analysis was used to identify 3 components (cPC1-3) that maximally separate neural activity across target locations. Right, quantification of the rotation in population trajectories (analysis as in Fig. 1L) predicted for an encoding model (left) compared to observed rotations in striatal (STR) and primary motor cortex (MOp). There were no significant differences between model and data. B Linear estimated dimensionality for the population of neurons recorded in each region. Each point is a separate dataset. **C** Random forest classifiers were used to decode the 4 trial types based on the neural peri-action responses with cross-validation and resampling to match the number of neurons across regions, i.e., same number of features (the product of the number of neurons and time bins) were used for interregional comparison. To estimate an upper bound of the classifier performance, classifiers were trained using all simultaneously recorded neurons (‘all’) across regions. **D** The Kalman filter was used to decode 3D hand trajectories based on the neural peri-movement responses with cross-validation and resampling to match the number of neurons across regions. Based on our finding that the linear mapping of neural responses to hand states may differ between reach versus pull phases, we trained separate classifiers for reach and pull phases and the decoded trajectories were concatenated. For simplicity trajectories on the Y-axis (outward) are displayed. See Supp. Fig. 4. for detailed decoding results. **E** Accuracy of decoded trajectories was evaluated using the coefficient of determination (R^2^) separately for reach (left) and pull (right) phases, i.e., we quantified how much variance in the actual hand trajectories was explained by the decoded trajectories during reach and pull phases. Top, Accuracy (R^2^) was compared between MOp and dSTR with the number of neurons matched by resampling, while the upper bound of accuracy was estimated by decoding from all MOp and dSTR neurons combined (black filled circles). Bottom, Decoding accuracies of the three resampled brain regions were compared.

While this argues broadly in favor of dSTR involvement in action specification, there is still a substantial diversity of accounts within this broad class that we sought to further refine. For example, one recent proposal (*15*) that could reconcile these results is that only the same “gross” action (*i.e*. reach-to-pull) is selected in dSTR whereas MOp determines fine variation in movement parameters (*i.e.* reach angle, pull force). We note that this would be surprising given that striatal activity is strongly correlated with continuous movement kinematics (Fig. 2A) (*44*, *55*, *56*), dSTR can be necessary for specification of movement parameters (*34*, *57*, *58*) and dSTR activity is sufficient to determine continuous kinematics (*32*, *56*). Nonetheless, these previous observations were generally made in tasks with single, highly stereotyped actions or highly variable movements rather than discrete selection between targets.

As with selection models broadly, a specification model in which subcortical dSTR activity only defines the gross action (shared across a large set of similar movements that only differ in the details of their kinematics and forces) whereas the motor cortex specifies fine parameters (*15*) makes several testable predictions. First, such a conceptualization implies that the dimensionality of population activity (*59*) should be *lower* in dSTR (one reach-to-pull action selected in all trials) when compared with MOp (four distinct reach-to-pull parameterizations specified) in this behavior. In contrast, if STR specifies movement parameters then the dimensionality should be the same or greater as compared to motor cortical activity. We found that striatal activity exhibited a *greater* linear dimensionality than activity in MOp during the reach-to-pull task (Fig. 2B; note: the nonlinear dimensionality is similar, but not lower in STR (*60*)).

Second, if it is the case that dSTR activity only determines the gross action and not fine parameter differences across conditions then our ability to classify trial type from dSTR population activity would be worse than classification using MOp ensemble activity (as implied by Fig. 5a (*28*); Fig. 6 in (*15*)). To evaluate this possibility, we trained decision-tree based nonlinear (random forest) classifiers to predict the trial type based on the activity of MOp, dSTR and MOs/ACA populations during movement execution (see Methods; pre-movement epochs were also examined with similar results). Trial types could be decoded with accuracy much higher than the chance level (25%) from all populations. We found no significant difference in the classification of trial types between MOp and dSTR populations across all datasets disconfirming the prediction of a gross selection model (Fig. 2C; Supplemental Fig. 4). Importantly, this classification performance is not ubiquitous. Activity in MOs/ACA, a premotor area critical to performance of this task, was nonetheless significantly worse at classification (Fig. 2C; One-way ANOVA; *F*_2,18_=5.44, *P*=0.014 with multiple comparison test *P*<0.05).

Third, a model in which striatum only selects gross action identity further implies that decoding of continuous movement kinematics from dSTR activity should be worse as compared to MOp. To address this question, we developed continuous time Kalman filter based decoders to reconstruct forelimb kinematics tracked with DeepLabCut (*61*) using MOp, ACA/MOs or STR neural activity (see Methods). We examined how the linear relationship between the activity of neurons and the state variables changed through the reach and pull phases.

Decoding accuracy was compared between neuron-number matched MOp and dSTR populations, which was also compared to the combined MOp+dSTR population that served as an approximate upper bound of decoder performance. Decoding accuracies did not significantly differ (One-way ANOVA; reach: *F*_2,18_=0.51, *P*=0.61; pull: *F*_2,21_=0.63, *P*=0.54). For the recording sessions with simultaneous recording of all three regions, we compared performance of MOp, dSTR, and MOs/ACA decoders. A significant main effect of the region was found (One-way ANOVA; reach: *F*_2,12_=17.06, *P*=0.0003; pull: *F*_2,12_=34.49, *P*=1.06×10^-5^), and the multiple comparison test with Bonferroni correction revealed comparable decoding accuracy between the neuron number matched MOp and dSTR populations (*P*>0.9), whereas a significantly lower accuracy was obtained from the MOs/ACA population (*P*<0.0005) despite MOs/ACA→STR activity being necessary for task performance (Supplementary Fig. 2).

### Single neuron action parameter encoding

The above analyses all support a model in which dSTR, like MOp, is critical for the specification of movement parameters, yet the population level analyses also suggest differences between the simultaneously recorded dSTR and MOp populations (e.g. linear dimensionality, Fig. 2B). In particular, one possibility is that dSTR exhibits preferential correlation for a subset of movement parameters. To look for such preferential encoding we next examined the discriminability of individual neuronal activity as a function of reach direction and pull load with a measure of discriminability (*d’*; see Methods). Briefly, *d’* was computed by first taking the difference between the mean firing rate of each trial type and that of the remaining trials. The products of differential firing rates and 2D vectors corresponding to each trial type were summed to yield *d’* scores across time within the 2D *d’* space. Individual units were classified into 8 classes based on which trial types allowed for significant discrimination (Examples shown in Fig. 3A and Supplemental Fig. 5).

**Figure 3.**
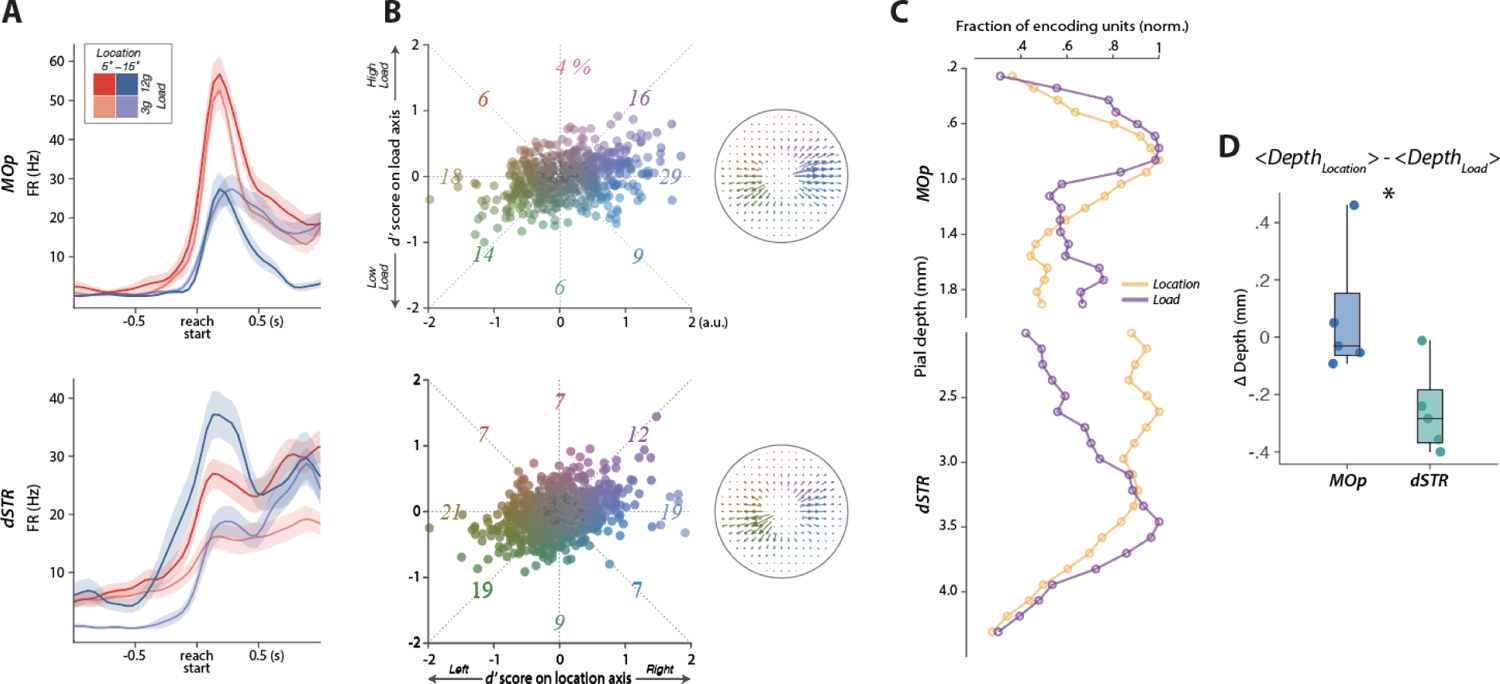
Quantifying MOp and dSTR single neuron encoding of trial conditions. **A** Example individual neuronal firing rates aligned to reach start (t=0) are plotted across the four trial types as mean and standard error (shaded). Top, A MOp unit preferentially responded when reaching to the joystick at 5°. *Bottom*, A dSTR unit further increased its firing rates when reaching for the joystick with a higher load. **B** Left, All MOp (Top) and dSTR (Bottom) single units were distributed in a 2D encoding space comprising joystick location (horizontal) and load (vertical) axes (See Methods for description of the encoding space and placement of single units). The 2D coordinates are each unit’s distance from the origin averaged across the 1-s post-reach start period. Units that passed a significance test using each unit’s trial-shuffled distribution, i.e., units with a significant encoding of movement parameters, were color coded based on their coordinates in the encoding space, while units that failed the significance test were marked as small gray circles. For example, the unit in **A** (top) that encodes 5° location but not the load is located in the mid-left part, while the unit in **A** (bottom) that encodes the higher load is located in the upper-center part of the space (See Supp. Fig. 5. for more example units). The color-coded numbers represent the percentage of encoding units classified into the 8 equi-distanced encoding groups. Right, Quiver plots were generated to summarize the distribution of units within the encoding space. Single units were spotted in the partitioned encoding space (21-by-21) and counted. The counts were normalized across brain regions for inter-regional compassion and translated into the length of arrowed vectors. **C** To compare the spatial distribution of encoding units across the corticostriatal depths the fraction of location- and load-encoding units were counted in depth bins and normalized to the maximum fraction across depths. MOp units encoding the two movement parameters display similar distribution concentrated around the depth bins within the cortical layer 5. In comparison, the dorsal and ventral dSTR subregions were preferentially populated with location-encoding and load-encoding units, respectively, displaying a higher degree of spatial segregation between the two encoding groups. **D** The spatial segregation of dSTR units encoding the two movement parameters observed in **C** led to a significant difference in Δ depths (difference in mean depths of the units that encode joystick location and load) between MOp and dSTR (n=5 mice).

Individual neurons in MOp distinguished both target location and pull load and were distributed across all sections of the *d’* space (Fig. 3A-B). A neuron for which *d’* trajectory surpasses the 95% confidence interval of the trial shuffled *d’* distribution (Methods) was considered to encode a task variable(s) with statistical significance. 67% of MOp neurons satisfied this criterion. Of these MOp neurons, a much greater fraction (47%) displayed a directional tuning, while fewer neurons (10%) displayed a load tuning (Fig. 3B). Among the directionally tuned MOp neurons a greater fraction (62%) was tuned for the contraversive (right) target (consistent with prior observations in primate MOp (*62*)). Interestingly, despite providing relatively poor decoding of trial-by-trial differences in movement kinematics, a similar fraction of individual neurons in ACA exhibited differential activity as a function of trial type (56%; Supplemental Fig. 5). Similarly, we found that striatal populations discriminated between parameter types and did not appear to preferentially encode direction (kinematics) nor force (kinetics). A substantial fraction (61%) of units showed significant discrimination in neural activity depending upon trial types with distinct kinematics or kinetics or conjunctions of the two (Fig. 3B).

### Broad spatiotemporal distribution of movement parameter encoding in striatum

Building upon anatomical evidence for topographic projections from cortex to striatum (*13*, *14*, *16*), many models propose that active neuronal ensembles for specific actions are restricted to tight spatial domains in STR (*28*). At the same time, action parameter specific correlates are distributed across corticostriatal projection classes (*44*); these classes have idiosyncratic and distributed anatomical projections to dSTR (*49*, *63*). This suggests that action parameter-specific ensembles could be more spatially distributed in striatum. To estimate these spatial distributions we next examined individual neuronal *d’* as a function of anatomical location.

First, we examined the spatial distribution of types - neurons with significant *d’* scores for either joystick load or target location parameters (Fig. 3C; for ‘mixed’ types see Supplemental Fig. 5). In MOp, we observed a relatively homogeneous distribution of types across the cortical layers with neurons with a significant *d’* concentrated in layer 5 for both types (Fig. 3C; Pearson correlation; all Rho values > 0.69; all *P* values < 0.001). Although we note that a modest shift in depth was apparent, putatively consistent with previous observations of preferential direction encoding in deep layer 5 (*44*). In contrast, we observed a more inhomogeneous spatial distribution in STR. We found neurons with preferential target location correlates were densely populated dorsally, whereas neurons with preferential joystick load correlates were found in densest numbers more ventrally. Accordingly, the Pearson correlation of neuron fractions that encode target location and load was reduced (Pearson correlation; Rho = 0.41; P = 0.07). While load- and location-encoding MOp units were located at comparable depths, the pull force encoding STR units were ventral to the reach direction-encoding units (delta depths MOp vs. dSTR; independent *t* test, *t_8_* = 2.68, P = 0.028; Fig. 3D).

We next examined when during execution of the reach-to-pull action *d’* tended to peak for different trial types. In MOp, normalized *d’* scores of these three encoding types displayed similar peri-action temporal patterns, as there was no significant interaction between time and encoding types (Supplemental Fig. 5; repeated-measures ANOVA; *F*_118,43837_=0.95, *P*=0.63). In contrast, we observed a significant temporal segregation in STR as the encoding of target location preceded the encoding of the joystick load (repeated-measures ANOVA; *F*_118,53336_=2.24, *P*=3.03×10^-13^). Thus, single neurons in STR provide similarly robust discrimination of movement parameters in roughly equal proportions to those in MOp; however, STR populations provide a less biased encoding that is also more distributed over space and time during reach-to-pull action.

### Cortex and striatum conjointly specify movement parameters

The above analyses provide additional evidence that dSTR activity, like that of MOp, is sufficient to classify actions and carry information about continuous kinematics of reach-to-pull actions. We found little evidence that the dSTR was categorically biased towards a subset of movement parameters (*9*, *15*, *30*, *31*, *33*). Moreover, there are many lines of causal evidence that basal ganglia output and dSTR activity in particular determines continuous kinematics of movements (*32*, *34*, *39*, *56*, *64*, *65*). Together these data point to a model in which dSTR output is critical for specification of movement parameters.

However, it remains in principle possible that dSTR correlates are inherited from MOp despite basal ganglia output playing little role in specifying fine parameters of movement execution. To quantitatively evaluate how trial by trial variations in observed MOp and STR activity account variation in reach angle and pull force we used a trial-based, linear committee decoder (*44*) that allows a direct comparison to models without having to consider continuous time dynamics of activity. Decoding performance using this approach was in general very good and could capture transitions around all switches in block conditions (Fig. 4A).

**Figure 4.**
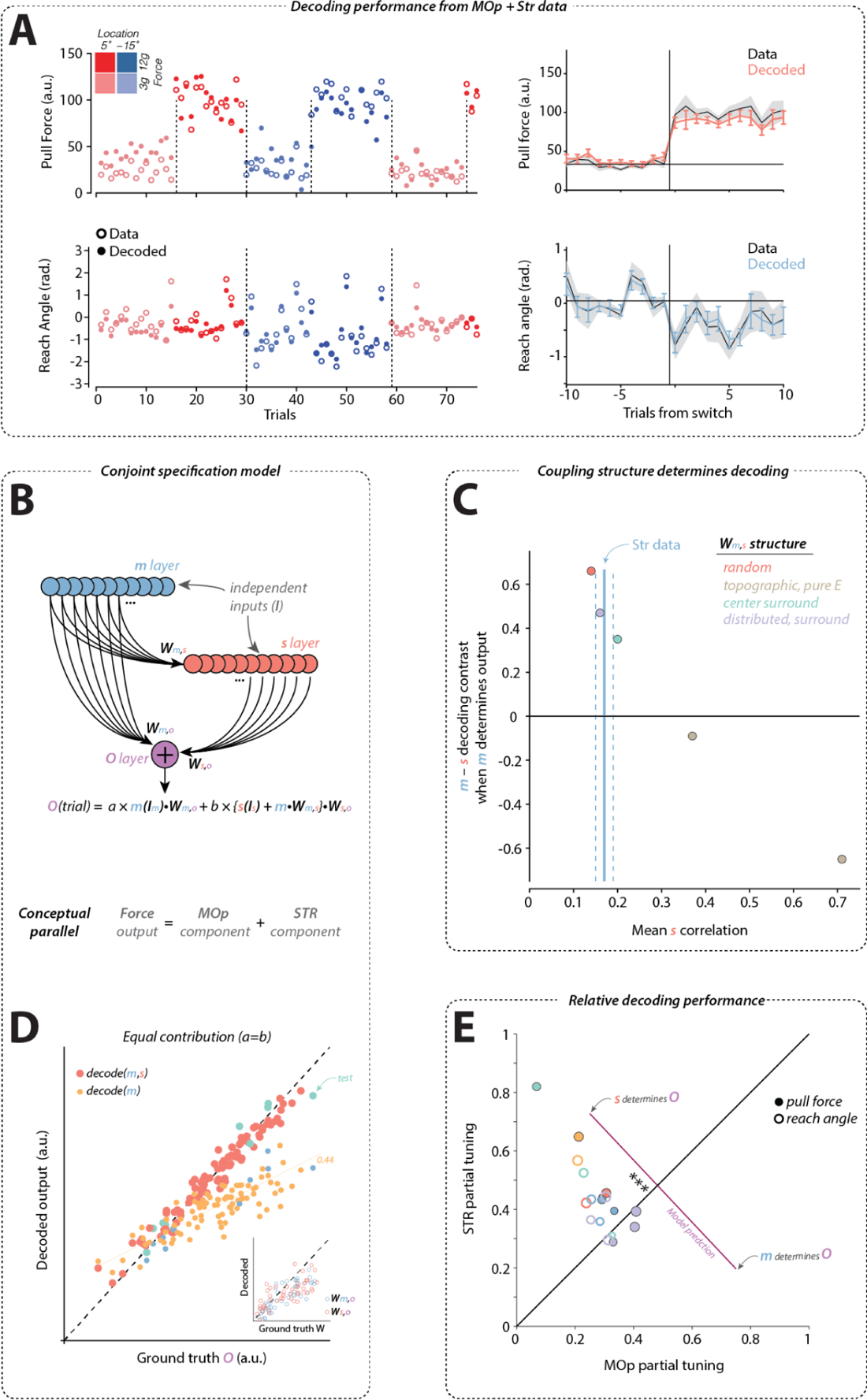
A conjoint specification model can best explain relative decoding performance of MOp and STR. **A** Observed (open circles) pull force (upper) and reach angle (lower) are plotted for each trial for 4 trial types (as indicated by color code). This was compared to predictions from a linear committee machine decoder (filled circles; see Methods; as in (*44*)). Right plots indicate the mean and standard error (shaded) for data (black) and the mean and standard error (bars) for decoded (colored) around all up-switches in pull force (upper) and all down switches in target angle (lower). **B** Schematic for two populations of units connected as in corticostriatal connectivity. ***m*** layer (blue) corresponds to MOp and ***s*** layer (red) corresponds to STR. The output, O, was modeled as a weighted sum of output from the ***m*** and ***s*** layers. **C** The difference in decoder tuning derived from ***m*** or ***s*** populations is plotted as a function of the pairwise correlation of activity in the ***s*** population across trials for a model configuration in which only ***m*** determines ***O***. Only decoders that recover positive values are accurately detecting model structure. Colors correspond to different formulations of ***W_m,s_*** structure (see Methods). Solid, vertical blue line is the mean pairwise correlation in Str data across all sessions (dashed lines indicate standard deviation). **D.** The same committee decoder performance as applied to encoding model data and plotted for decoded predictions (*y-axis*) compared to observed ground truth output (*x-axis*) for a decoded using the full ***m*** and ***s*** layer data (larger, redder circles) and partial decoding from just ***m*** layer activity (smaller, yellow circles). Inset, the inferred weights from the committee decoder are well correlated with the ground truth ***W*** (*x-axis*). **E.** Decoding of pull force (filled) and reach angle (open) from data from Str (*y-axis*) is plotted against MOp (*x-axis*) for all sessions. Purple line indicates the expected trade off in decoder performance for models where either exclusively ***s*** or exclusively ***m*** determines ***O***. Significance testing revealed a greater contribution from STR compared to MOp; *signrank test; ***, p<0.001*).

**Figure 5.**
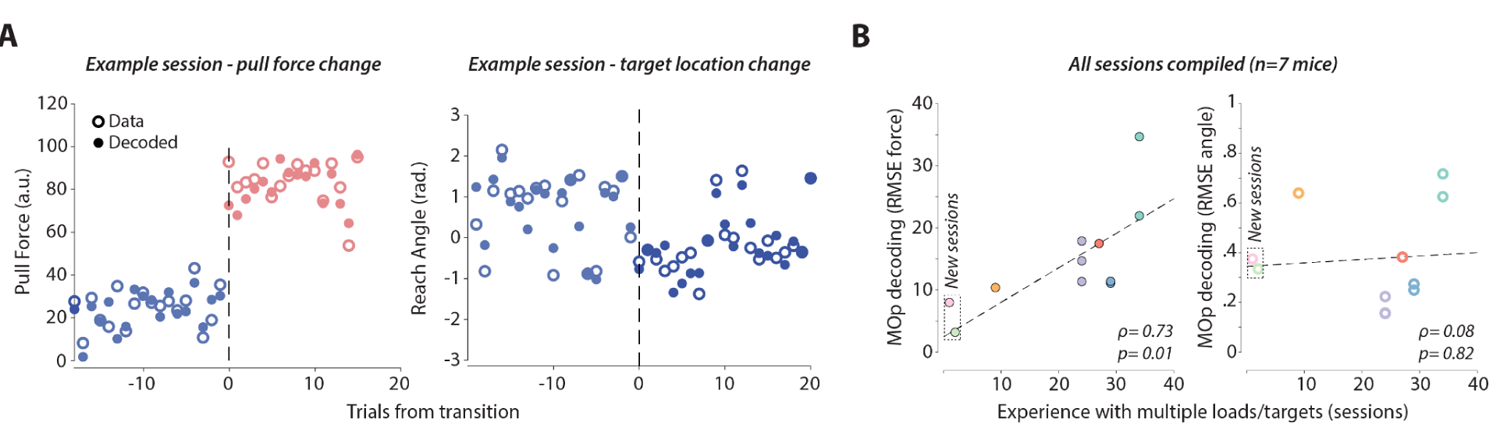
Encoding of novel movement parameters may reflect MOp-driven specification of novel forces. **A** Observed (open circles) pull force (left) and reach angle (right) are plotted around the transition to a new pull load and target location in an example session. Decoding from MOp population recording as in Fig. 4. **B** Quantification of the accuracy of MOp-alone decoder performance for applied force (left) and reach angle (right) plotted as a function of the total number of sessions of experience with multiple targets/loads. In addition to previous mice (N=5) two new mice were trained on a single joystick load and single location. We then compute the decoding performance on the first day as they transitioned to a novel load and target location (‘New sessions’). Regression line is fit across all data (N=7 mice; 11 sessions). Significant correlation was also detected in just the subset of trained mice (N=5; p<0.05, not plotted) for force. Colors indicate individual mice.

If two independent populations of units conjointly determine output, a decoder trained on both populations will correctly recover that each population contributes 50% to the predicted output (Supplemental Fig. 6). Inspired by the organization of corticostriatal circuits (*6*), we next consider a structure in which a population akin to MOp (‘**m**’) provides input to a downstream population akin to STR (‘**s**’; via **W_m,s_**; but not *vice versa*) and a weighted sum of the outputs of both populations determines the observed output (‘**O**’) akin to a continuous variation in a movement parameter (Fig. 4B; see Methods for details) (*16–18*). At one extreme, when **W_m,s_** is unstructured (random) and output is purely driven by **m** the decoding is appropriately much better from **m** compared with **s** (Fig. 4C; red point). At the other extreme, when **W_m,s_** acts to pool and denoise **m** activity but the output is determined by **m** only, then decoding can indeed indicate a greater contribution from **s** than **m** that is not present (Fig. 4C; yellow point). Consistent with the interpretation that this is due to pooling and denoising, this effect is well predicted by an enhanced pairwise correlation of **s** activity (Fig. 4C). There is little empirical support for a purely excitatory and focused topography with no lateral inhibition in corticostriatal connectivity (*16–18*, *66*) and similarly we observed *reduced* rather than elevated pairwise correlations in dSTR activity patterns (STR: 0.17±0.02 (s.d.); MOp: 0.19±0.03; Supplemental Fig. 6). Distributed topographic connectivity combined with modest lateral inhibition, consistent with corticostriatal anatomy (*16–18*, *66*), was able to recapitulate both reduced pairwise correlations in **s** and decoding performance that reflected ground truth contribution to output (Fig. 4D-E).

We quantitatively evaluated decoding performance in our datasets using the same decoding methodology. Across all recording sessions we found similar, good performance of decoders trained on MOp and STR populations (Fig. 4D). Plotting decoding performance in STR vs MOp revealed a negatively sloped trade-off in good quantitative agreement with model predictions over a range of roughly equal contribution (Fig. 4E). The data was skewed towards models in which MOp and STR make relatively balanced contributions. Specifically, we observe reliably better decoding from STR activity for both pull force (p=0.008; *ranksum* test) and reach angle (p=0.0005; *ranksum* test). The observed high decoding performance from STR and worse decoding from MOp when both areas are used jointly in decoding is thus more consistent with predictions of a model in which the conjoint activity of MOp and STR determine movement parameters.

### Neocortical contribution to movement kinetics decreases as experience of multiple movement parameters

The majority of prior theorizing about cortico-basal ganglia function has argued for dissociable functional roles for cortex and subcortical striatum. Our data here are best explained by a model in which cortex and striatum conjointly specify the parameters of dexterous reach-to-pull actions. However, we found the relative STR contribution appeared to vary across subjects as indicated by the relative contribution of dSTR and MOp decoders being distributed along the line defined by varying relative contribution across **m** and **s** layers (purple line in Figure 4D).

On the one hand, this scheme seems to diverge from dissociable functional roles and may even seem wasteful to use two forebrain areas in the same function of specifying movement execution parameters. On the other hand, we may ask what advantage is afforded by conjoint specification across multiple areas. A key aspect of the adaptive and flexible control of voluntary action is the ability to adjust the parameters of action to achieve goals based upon the current context, past experience, behavioral state, competition, etc. From this perspective, conjoint specification could allow more diverse or even independent mechanisms to modify the same action. For example, rapid top-down explicit control (say following instruction (*2*)) could modify movement speed via direct modulation of MOp activity. Alternatively, experience with selective reinforcement for actions performed within a specific range of parameters (*56*) could implicitly modify movement via dopamine-dependent modulation of synaptic weights on STR neurons (*67*) without modifying MOp population activity. In the case of consistent well-learned actions we might expect specification of action parameters by STR to be particularly critical given ample opportunity to learn implicit modifications to movements (*31*). Recent work studying stereotyped forelimb movements found that MOp is only critical for acquisition and STR critical after initial learning (*68*, *69*). This would suggest that when learning a new action parameterization MOp might likewise be particularly critical with STR coming to contribute only after substantial experience (*31*).

To examine this possibility, we trained a new pair of mice on a single target location and joystick load. We then recorded neural activity primarily in MOp during the very early exposure (<3 sessions prior) to a novel target location and/or joystick load requirement. To compare MOp decoding performance we examined reconstruction error (RMSE). We find that in these new mice experiencing a novel joystick load, decoding performance from MOp activity was better than observed in any of the 9 datasets examined in well trained animals (Fig. 5). Indeed, in well trained mice MOp decoding performance was significantly negatively correlated with the extent of training experience and the new session data were consistent with that trend (Fig. 5B). However, we found no such correlation for reach angle decoding suggesting a sustained, critical role of MOp even with extensive training.

## Discussion

Canonical models invoke an abstracted representation of action in dSTR subserving selection (*15*) and initiation (*13*, *14*) of gross actions in distinction from the continuous specification of movement kinematics and kinetics involved in online control of execution by MOp (*5*, *10*). In contrast, recent proposals highlight critical roles of dSTR in determining the detailed movement parameters underlying goal-directed actions (*6*, *30–32*). As we show using quantitative, but simple models, existing datasets are equivocal on this distinction (Fig. 1). To distinguish between these models requires studying MOp and STR activity as an animal chooses between the same action with modest variation in the parameters of the underlying movements - a challenge for rodent studies. Here we developed such a paradigm and our data are uniquely consistent with a model in which the fine movement parameters underlying a reach-to-pull action are the product of conjoint specification by MOp and STR (*6*, *30*, *70*). Moreover, our data are inconsistent with models invoking a circumscribed selection of ‘gross’ actions by STR (*15*, *38*, *71*).

This work replicates and builds upon previous work that has described robust dSTR encoding of kinematics in the case of both highly variable (*34*, *35*, *44*, *55*, *56*) and stereotyped (*32*, *72*) actions. Calcium imaging data has, in some cases, suggested an absence of dSTR encoding of movement kinematics (*36*, *37*). However, this may be a limitation inherent to calcium imaging in STR (*73*, *74*) or failing to examine variation in relevant movement kinematics in detail (*39*) as other imaging studies do find clear kinematic correlates (*75*, *76*). In the cortex, where direct comparisons between electrophysiology and imaging have been made, it is clear that calcium imaging can make fine behavioral correlates noisier and harder to identify (*44*, *77*). Moreover, we observed encoding of movement parameters to be distributed over millimeters of depth in STR. Imaging in mice has typically explored only a single depth plane (e.g. (*37*)) and thus may have missed anatomically distributed correlates. Finally, we show that activity in the putative premotor region in MOs/ACA - despite being critical for reach-to-pull actions - has significantly worse decoding of fine movement parameters. Thus, while non-negligible encoding of gross movement parameters may be ubiquitous (66, 67), fine (in distinction from “gross”) parameter variation of an action can be preferentially encoded in specific forebrain regions such as forelimb MOp and dorsal STR.

How might conjoint specification of movement parameters by MOp and STR be useful for different aspects of learned, adaptive voluntary action? What may seem redundant for one aspect of behavior (*e.g.,* executing a well learned action), may afford unique opportunities for other aspects of behavior (*e.g.,* differential learning mechanisms). Computational modeling with artificial neural networks demonstrates that distinct architectures (*e.g.* dense vs. low-rank connectivity) can achieve equivalent output performance for learnt actions (*60*, *78–80*). Consistent with this general insight from modeling, here we observed similarly good decoding performance from either MOp or STR activity (as in highly variable actions (*44*)) yet greater spatiotemporal separation of individual neuron encoding subcortically. STR receives dense innervation from MOs/ACA partially overlapping and partially distinct from MOp innervation (*17*, *18*, *21*, *50*, *81*) which may help to explain greater spatiotemporal distribution of movement variable encoding. MOs/ACA projections to STR appear critical for initiation of actions and exhibits the earliest pre-reach modulation of activity in mice (here) as described extensively in primates (*7*). MOs/ACA also projects densely to non-STR targets (*e.g.* zona incerta (*18*, *82*)) not as strongly innervated by MOp output. These extra-telencephalic projections may help to explain a critical role of MOs/ACA in the initiation of an action rather than or in addition to MOs/ACA→STR collateral projections (*83*). However, it is also the case that an extreme dorsomedial aspect of dSTR receives the densest ACA projections (*50*) and future work will be required to investigate a potential role of this more circumscribed subregion in action initiation.

Conjoint specification of fine movement parameters by cortical and subcortical brain areas may allow for distinct circuit mechanisms of learning and adaptation. There are many differences between afferent inputs to MOp and dSTR. For example, dSTR receives dense innervation from midbrain dopamine neurons, amygdala and intralaminar thalamic areas that innervate MOp less. The involvement of basal ganglia (*19*, *25*) and amygdala (*84*, *85*) in reinforcement learning has led to the proposal that the representations of movement parameters in dSTR is critical for slow learning of implicit movement specification (*31*, *67*). More rapid learning, in some cases involving explicit specification of movement parameters (*86*), is thought to be dependent on neocortex (*87*). Consistent with this possibility, we provide evidence across seven mice that lesser training extent, including two mice learning a novel pull force, is associated with an enhanced MOp contribution that slowly reduces with more extended experience of multiple required movement parameterizations (*69*). However, even in the well trained condition many lines of evidence indicate that MOp and STR continue to play a conjoint role in specifying the movement parameters underlying skilled forelimb movements in mice and mammals generally (e.g. (*10*, *44*, *56*, *87*, *88*)). The robustness to MOp lesion for some stereotyped (although not flexible) forelimb actions may reflect a differential capacity to compensate for loss of MOp function that arises with training (requiring >1 week (*89*, *90*)), rather than necessarily implying a sequential, functional hand off (“tutoring” (*30*, *32*)) between cortical and subcortical areas.

There has been some progress articulating putative learning rules in MOp (*91*) and STR (*56*) that might allow the flexible specification of fine parameters of goal-directed, skilled actions. However, many critical questions remain from the perspective of a conjoint specification model as proposed here. How are plastic changes across these cortical and subcortical circuit components coordinated? How is specificity across multiple distinct learned actions maintained? Why did conjoint specification of movement parameters in cortical and subcortical motor regions evolve? Perhaps it reflects a necessary hand off during development for the more slowly developing corticospinal innervation. These and other questions will be critical to address in future studies.

## Methods

All handling of animals and procedures were performed in strict accordance with the Janelia Research Campus Institutional Animal Care and Use Committee (IACUC) and the standards of the Association for Assessment and Accreditation of Laboratory Animal Care (AAALAC). Male and female mice, typically aged 8 to 16 weeks at time of surgery, were used in this study. Mice were water restricted (1 to 1.5 ml of water/day) with daily health checks. Water restriction was eased if mice fell below 75% of their original body weight. Slc17a7-Cre (VGLUT1-Cre) mice were generated by the Janelia Research Campus Gene Targeting and Transgenics Facility. Behavioral and neural data were taken from a total of 5 Slc17a7-Cre mice.

### A novel joystick apparatus

The joystick made of stainless steel was connected to two stepper motors that provided rotation in the pitch and yaw. A stepper motor rotary encoder tracked the joystick-attached stepper motor from which the joystick kinematics were inferred. The joystick could be robotically positioned to one of two yaw locations on an equidistant arc in front of the head-fixed mouse. The joystick was then rotated into a pitch angle that was within the grasp of a head-fixed mouse. An auditory cue was delivered indicating when the joystick was in position and the task of the mouse was to reach out to the correct yaw target location, grab the joystick, and pull along the pitch axis past a specified threshold (5mm). The pitch motor could be used to set a variable load by controlling the amount of current to the motor windings. The pitch motor’s holding load, which is the amount of force needed in order to move the motor one full step when the windings are energized but the rotor is stationary, was calibrated with a force gauge and translated to the force required to move the joystick. In these experiments we selected a 3g or 12g requirement.

Each trial began with positioning of the joystick and ended with a successful pull (> 5mm). Trials with incorrect responses (e.g., pushing the joystick past a threshold, 5mm) or timeout (the lack of pull or push for 10 s) were marked as unsuccessful. After the end of each trial the joystick was retracted away from the mice and returned to the home position to be placed back to the set position for the next trial after the inter-trial interval (7 s).

The design files for constructing the joystick hardware can also be downloaded from: https://www.dropbox.com/scl/fo/y4ozvuk3mgx61g19stthj/h?rlkey=s6soot4oazr9520nij5pgt1lx&dl=0

All joystick operations according to the user-defined task structure were programmatically controlled using a custom-written open source python package: https://github.com/janelia-pypi/mouse_joystick_interface_python.

Additional details and other previously published hardware and software and analysis code will all be found and maintained at: https://dudlab.notion.site/Conjoint-specification-of-action-in-neocortex-and-striatum-cebef9fc306947139f0ee70ce2dadeaa?pvs=4

### Training

Mice were trained for 3-7 weeks approximately 1 h every day before the initial data acquisition. For the first few days mice were acclimated to head fixation and the water spout that delivered drops of sweetened water. Once habituated with head fixation and water consumption the joystick was initially positioned in close proximity (5mm) to the resting right hand of the mice to facilitate discovery of the joystick with spontaneous limb movements. In the early phase of training, the pull threshold (<2mm) and load (<1g) were set to be low such that a slight pull triggered an immediate water reward. As mice learned to reach and pull the joystick, we gradually adjusted the task parameters - the initial position of the joystick (5 to 20mm from the resting hand), pull threshold (2 to 5mm), pull load (1 to 12g), and the delay for water delivery (0 to 1s).

When the mice achieved an expert level performance (>150 successful trials in 60 min) with the adjusted parameters, we introduced the block design, in which mice performed two repetitions of the four joystick location (Left; 5°, Right; −15°) and pull load (Low; 3g, High; 12g) pairs - 1) 5°/3g, 2) 5°/12g, 3) −15°/3g, 4) −15°/12g. All behavioral and neural data were taken when mice achieved an expert level with the varying task parameters.

### Silencing of MOp→STR or MOs→STR projection neurons (Supplemental Figure 2)

To examine the behavioral effect of silencing MOp→STR or MOs→STR projection neurons, rAAV2retro-hSyn-SIO-stGtACR2-KV-eGFP [3.0 × 10^12^ genome copies (GC)/ml] was injected to the dorsal striatum bilaterally (relative to lambda: 0.5 mm anterior; ±1.7 mm lateral; 2.8, 2.6, 2.4 mm deep; 30 nl per depth), labeling neurons projecting to the dorsal striatum. Viruses were obtained from Janelia Viral Tools (http://www.janelia.org/support-team/viral-tools). In randomly selected silencing trials a single pulse 473-nm bilateral laser was triggered during the inter-trial interval and terminated either by a trial completion or after 4 seconds.

### Extracellular electrophysiological recording

Before recordings, a craniotomy was made over the recording sites (MOp: 0.5 mm anterior, −1.7 mm lateral; MOs/ACA: 1.0 mm anterior, −0.3 mm lateral relative to bregma) at least 12 hours before recording under isoflurane anesthesia. All recordings were taken from the left hemisphere contralateral to the right forelimb that mice used to perform the task. The probes were centered above the craniotomies and lowered with ∼10 degree angle from the axis perpendicular to the skull surface at a speed of 0.2 mm/min. The tip of the MOp/STR Neuropixels probe was located at ∼4.2 mm ventral, while the MOs/ACA probe was lowered ∼1.5 mm ventral from the pial surface. Before each insertion surface of each probe was coated with CM-DiI (Invitrogen), a read fixable lipophilic dye, for histological verification of probe tracks. Exposed brain tissue was kept moist with phosphate-buffered saline (PBS) at all times, and craniotomy sites were covered with Kwik-Sil elastomer (World Precision Instruments) outside of the recording session. All recordings were made with open-source software SpikeGLX (http://billkarsh.github.io/SpikeGLX/). Spike sorting was performed using the Kilosort2 (https://github.com/MouseLand/Kilosort) template matching and clustering algorithms with manual curation of the detected spikes using Phy (https://github.com/cortex-lab/phy).

### Characterization of individual neuronal encoding of trial types with d’ (Figure 3)

To quantify the degree to which the 2×2 trial types of joystick loads (low vs. high) and positions (left vs. right) are encoded by individual neuronal firing rates, the discriminability index (*d’*) was computed for each trial type. The individual neuronal mean firing rate of each trial type was subtracted by the mean of the rest of the trial types with normalization by the sum of standard deviations.

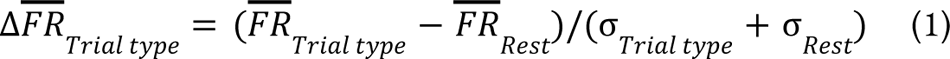

Applying eq (1) for each of the four trial types yielded four ΔFR values that were projected onto four vectors [1 1]^T^, [1, − 1]^T^, [− 1, − 1]^T^, [− 1, 1]^T^. These vectors correspond to the four location-load pairs, namely, right-high, right-low, left-low, and left-high, respectively. The sum of projected vectors yields 2D coordinates on an ‘encoding space’ that characterizes the neuronal encoding of the joystick location (horizontal axis) and required force (vertical axis). The length of this resultant vector was taken as the neuronal trial-type discriminability score, *d’*. Continuous time *d’* scores (trajectory) was computed for each 20-ms bin spanning −1 to 1 s relative to the reach start. *d’* captured the preferential tuning property of individual neurons and its visualization within the 2D *d’* space.

For a statistical significance test, trial-shuffled *d’* trajectories were obtained with 1000 shuffles, and their maximum distances from the origin (0, 0) were calculated. The 95% confidence interval was estimated from the maximum distance distribution (*mean* + 2 · *std*). If a neuron’s actual *d’* trajectory (i.e., without trial shuffling) had crossing(s) of this 95% confidence interval in one or more time bin(s), the neuron was considered to encode a task variable. The neuronal encoding was further characterized by subdividing the 2D encoding space by the eight equi-spaced vectors (Fig SXX). Single units were classified into the 8 classes by spotting which of the 8 equi-spaced vectors its max *d’* coordinates are best aligned to.

### Neural decoding of 3D hand trajectories with Kalman filter (Figure 2)

Our Kalman filter for neural decoding consisted of the state model and the observation model (*92*). The state model captures the linear function and its uncertainty, based on which the state variables (3D hand trajectories) evolve from time *t-1* to *t* with 20-ms timesteps. The linear relationship between the state variables and the activity of neurons is captured by the observation model. Parameters specifying the state and observation models that are jointly Gaussian were learned in the training phase. In the test phase, the held-out hand trajectories were estimated recursively through one-step prediction and measurement update. The model performance was evaluated as the coefficient of determination (R^2^) between the actual and estimated trajectories.

The state model is defined as:

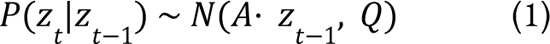

where *Z* is a three dimensional vector containing state variables (3D hand coordinates) per timestep (20 ms). *A* is a matrix that describes transition of state variables between timesteps. *Q* is the covariance matrix that characterizes the uncertainty around the estimated state transition. The initial hand coordinates *Z* was estimated as the sample mean and covariance from the training dataset.

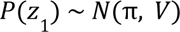

The observation model is defined as:

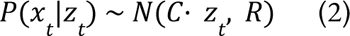

where *X* is a *n*-dimensional vector (*n* = the number of neurons) containing the spike count of neurons per timestep. *C* is a matrix that captures the linear mapping between the state and observation variables. R is the covariance matrix that characterizes the uncertainty around the estimated observation variables. The model parameters θ = {*A*, *Q*, π, *V*, *C*, R} were estimated analytically on the training dataset during the training phase. In the test phase, the 3D hand trajectories of the held-out trials were estimated recursively following one-step prediction and measurement update.

One-step prediction:

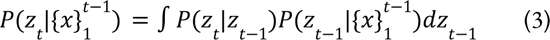

Measurement update:

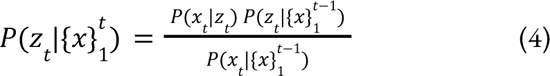

A leave-one-out cross-validation was used to train and evaluate the model from separate datasets. When there are two (MOp, dSTR) or more (MOp, dSTR, MOs/ACA) simultaneously recorded populations, we trained and tested the decoder with or without resampling the neurons from each population. The purpose of resampling was to match the number of neurons when comparing the decoder performance between populations. Resampling was repeated for a total of 30 iterations. Models trained with all neurons without resampling provided an approximate upper limit of decoder accuracy.

### Dimensionality estimate (Figure 2)

Linear dimensionality was estimated as described previously using principal component analysis (*60*). We report the max dimensionality across recorded datasets; however, the difference between STR and MOp were robust across a broad range of resampled population sizes using 10-fold resampling.

### Random forest classifier (Figure 2)

To compare discriminative encoding of trial types (four location and load combinations) across neural populations, we trained decision tree-based random forest classifiers via the scikit-learn’s RandomForestClassifier implementation. Binned spike count data were organized in 3D matrices whose dimensions were the number of neurons, the number of time bins, and the number of trials. Only data from trials with successful reach-to-grasp performance were used to train and test classifiers. Stratified 5-fold cross validation (scikit-learn) was used to train and test classifiers with balanced distribution of the four trial types within each fold. For inter-regional comparison of decoding accuracy the number of neurons were matched across simultaneously recorded neural populations by random sampling over 100 times. Data points shown in Fig. 2C each represents the median decoding accuracy of the 100 accuracy values calculated for each resampled data. Python codes used for training and testing random forest classifiers can be found here: https://github.com/jup36/MatlabNeuralDataPipeline/tree/master/neural_encoding_trial_types_js2p0

### Committee decoder design (Figures 4 & 5)

To assess the contribution of distinct neural populations to forelimb movement, we used a linear decoder to estimate the trial by trial parameters of reach angle and pull force from integrated neural activity (windows:) during the reach and pull phases, respectively. The decoded estimates were then correlated with the actual joystick trajectories. The decoder seeks to identify an optimal (minimization of least squares) linear mapping (*W_decode_; dimension Neurons x 1*) between the neural population activity (*R; dimension Trials x Neurons*) and the chosen behavioral parameters (max pull force or reach angle; *B dimensions Trials x 1*):

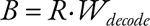

We solve for *W_decode_* as the mean of the estimates *W* computed (using *psuedoinverse*(activity) * data) from random batches of training trials (batch size: 75% population; batch number: 200-500; model: batch size: 30; batch num: 200). For cross validation, we evaluate the decoder on total trials or held out trials to look for good generalization of *W_decode_* performance.

Based upon ground truth models (see below) we found the best evaluation of decoder performance to be the slope of the relationship between predicted and observed data for reach angle and pull force separately (example in Supp. Fig. 6 and Fig. 4). For evaluation of decoding quality from a given population we use root mean squared error (RMSE; Fig. 5). Estimates of average pairwise correlations are obtained by taking the mean of correlations in the upper triangular portion of correlation matrix of integrated activity per trial.

### Encoding simulated data (Figure 1)

To produce the activity dynamics simulated in Figure 1 and Supplemental Figure 1 we generated simulated networks (typical N=120 units) in which activity profiles were produced as a combination of random lags at the typical timing of reach onset or pull onset, variable durations, and an independent response timed to the latency of reward delivery (gain: Poisson random lag: λ=3). Individual profiles were generated as a combination of a half Gaussian rise (median: 45; uniform random: [25:65] & median: 80; range: [25:300]) and a half Gaussian decay (median: 150 & median: 200). Lags were determined as a single sample per unit from a Poisson random distribution determined by λ=1:120 and λ=9:9:1080. Tuning to reach angle and pull force were generated from cosine tunings and uniform random monotonic gain tuning (uniform random, range=[0:0.8]), respectively (as in canonical motor direction tuning analysis (*40*) and prior observations in mice, e.g. (*34*)). Observations reported are consistent across a wide range of parameter values.

### Two population decoding simulated data (Figure 4)

To produce the activity dynamics simulated in Figure 4 we simulated the total spike count per trial for N=100 trials. To produce tuned activity we considered inputs, independent across the two populations termed **m** and **s**. Each unit received noisy input tuned across a range of monotonic gains - uniform distributed between [-0.15 and 0.85] - and a normally distributed source of noise N(0,σ=1). 15% of the coupling between m and s layers is determined by the matrix W_m,s_. We considered several different structures. With the exception of the random W_m,s_ matrix (uniform: [-0.5:0.5]) all matrices were produced by a Gaussian distributed weights with an excitatory and inhibitory component uniformly shifted by one unit across the population. Gaussian excitatory and inhibitory kernels had the following dispersion parameterizations:

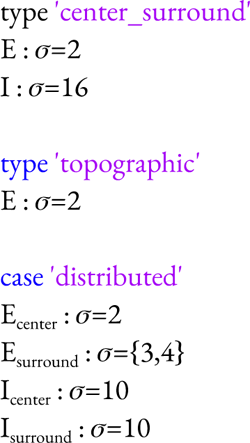

Example code for simulations will be made available at: https://dudlab.notion.site/Conjoint-specification-of-action-in-neocortex-and-striatum-cebef9fc306947139f0ee70ce2dadeaa?pvs=4

## Acknowledgments

We thank all the members of the Dudman and Gallego labs for project feedback; J. Arnold, Janelia Experimental Technologies designed the hardware for the joystick apparatus and P. Polidoro designed the software control both under extensive interaction and iteration with J. Park; J. Keller provided technical assistance for several hardware issues; the Janelia Viral Resources team for supplying reagents; M. Copeland and B. Foster for assistance with histology. The analysis in Figure 5B was inspired by a helpful discussion with Emmett Thompson (SWC) and Francesca Greenstreet (SWC) at iBAGS 23. This work was supported by Howard Hughes Medical Institute where J.T.D. is a Senior Group Leader at the Janelia Research Campus.

## Author contributions

Conceptualization: JTD, JP, PP, JA Methodology: JP, PP, JA, JAG, JTD

Investigation: JP, CF, JTD Visualization: JP, CF, JTD

Funding acquisition: JTD

Project administration: JTD

Supervision: JTD, JAG

Writing – original draft: JP, JTD

Writing – review & editing: JP, JTD, JAG, BM, CF, JA, PP

## Competing interests

Authors declare that they have no competing interests.

## Data and materials availability

All data, code, and materials used in the analysis will be made available on public databases as described in methods (code: GitHub; data: FigShare) and collected together at https://dudlab.notion.site/Conjoint-specification-of-action-in-neocortex-and-striatum-cebef9fc306947139f0ee70ce2dadeaa?pvs=4

Supplementary Materials:

## Supplemental figures

**Supp. Fig. 1.**
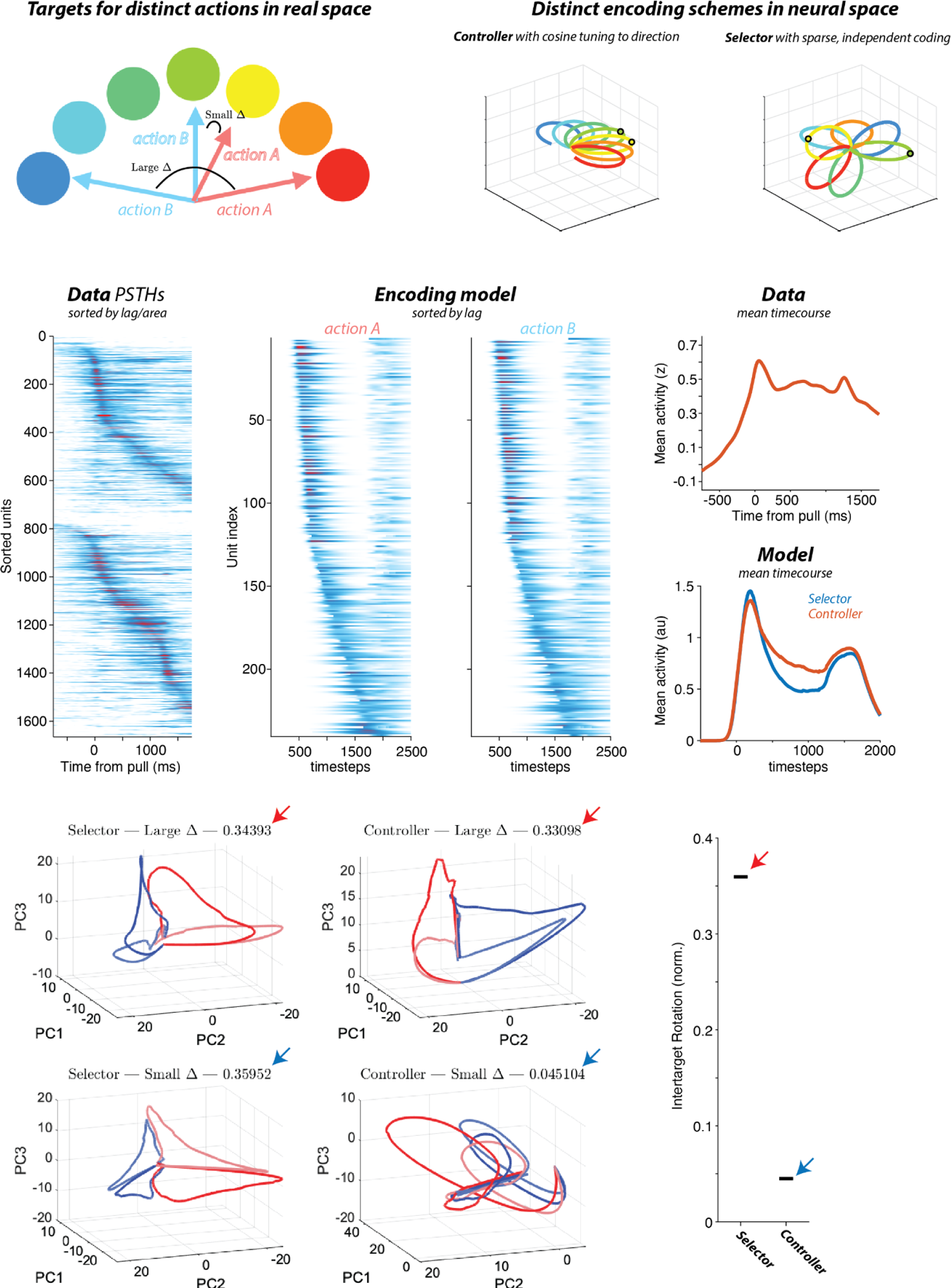
Elaboration on the encoding model A Consider a set of different targets which two actions. **(A & B)** may target, e.g. a reaching movement of the forelimb. We are interested in comparing the expected neural activity differences between two actions as a function of whether those actions are similar in real space (small intertarget angle) as opposed to far apart. **B** we consider two different kinds of encoding schemes in the space of neural activity dynamics (3 dimensions illustrated; color code as in A). In the case of a “controller” model (left) neural activity mimics the real space of different actions - e.g. cosine tuning of activity to reach angle laying on a flat manifold in neural space). The power of sparse coding scheme useful for a “selector” model is that activity is projected in separate directions for any given action in the space. This allows all actions to be similarly far in their encoding in neural space (i.e. a sparse code). Thus, for similar actions in a controller model the neural activity will be nearby (left, black line) whereas for a selector model even similar actions will have distant representations (right, long black line). **C**, neural activity in both cortex (upper plot) and striatum (lower plot) has complex dynamics of activity with neurons tuned to many different lags and moments of the reach-to-grasp performance. **D**, we simulated the selector and controller model (described in methods) for two similar actions. PSTHs are aligned to highlight the dominant temporal structure with harder to observe differences as a function of action A and B. **E**, The mean response profile over time for all units. **F**, the meant response profile averaged over all units in the encoding models. **G**, illustrative population dynamics projected onto the leading principal components (PC1-PC3) for the controller and selector models for actions with a large angle and those with a small angle (plots as labeled). Numbers indicate the normalized Procrustes rotation required to map activity from action A onto action B. H, note that only the controller model predicts small rotations for similar actions. See text and Methods for additional details.

**Supp. Fig. 2.**
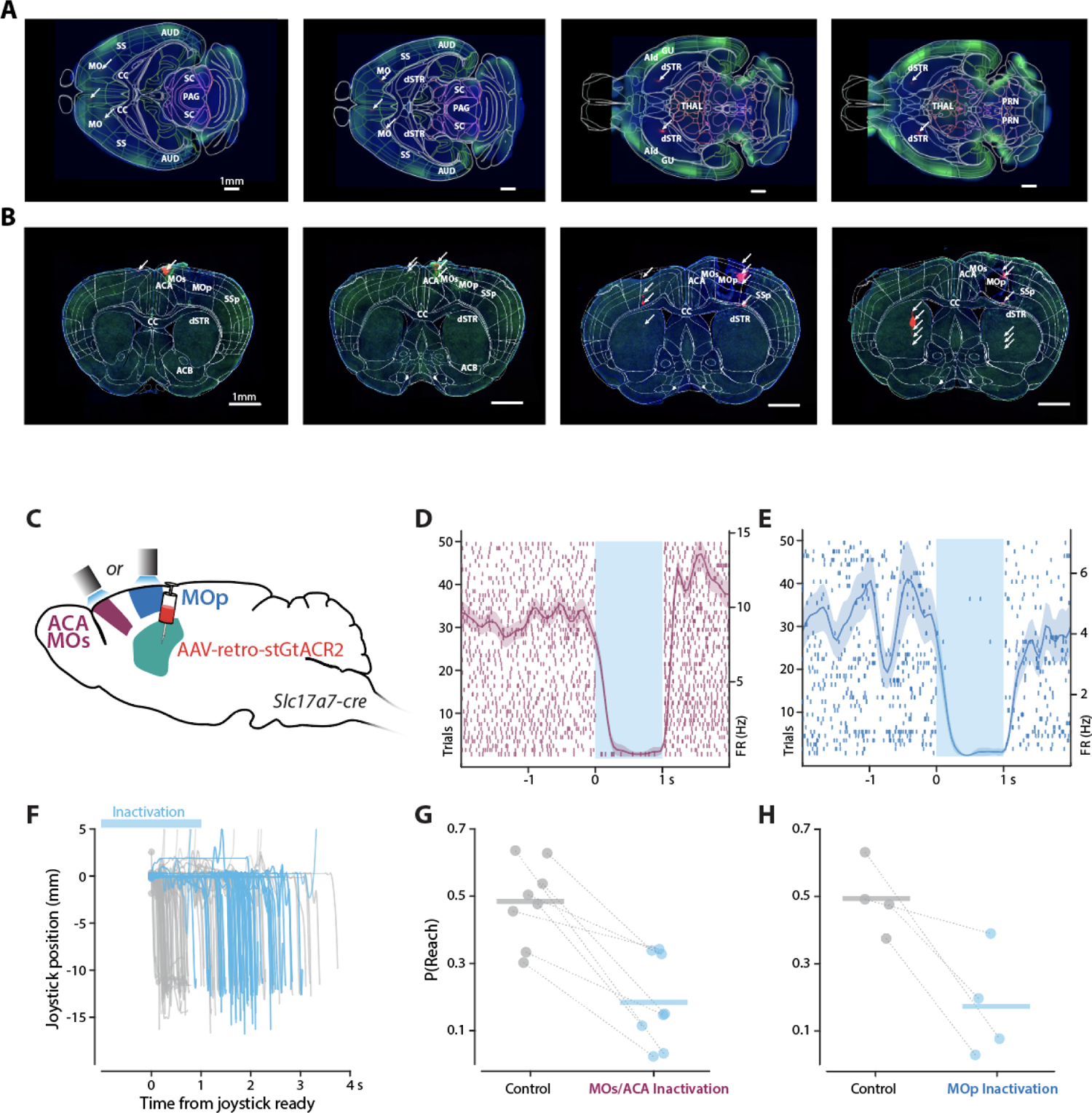
Histological verification of electrophysiological and silencing experiments. **A** Histological reconstruction of the probe track labeled with DiI (highlighted in red, marked with white arrows) superimposed with DAPI (in blue). The presence of eGFP (in green) signifies the expression of GtACR2 for optogenetic silencing. Four exemplary horizontal sections are presented in order from dorsal (left) to ventral (right). **B** Histological reconstruction in representative coronal sections arranged from anterior (left) to posterior (right). The color scheme matches that of A. **C** To inactivate ACA/MOs or MOp neurons projecting to dSTR GtACR2 was expressed with rAAV2-retro in Slc17a7-Cre mice. **D** An example ACA/MOs that exhibited robust silencing during a 473-nm laser. **E** An example MOp neuronal silencing. **F** Inactivation of either ACA/MOs or MOp significantly reduced reach-to-grab occurrences as exemplified by joystick trajectories with (in blue) or without (in gray) the optical silencing. G-H A significant reduction in P(Reach) was observed during inactivation of ACA/MOs (G) or MOp (H).

**Supp. Fig. 3.**
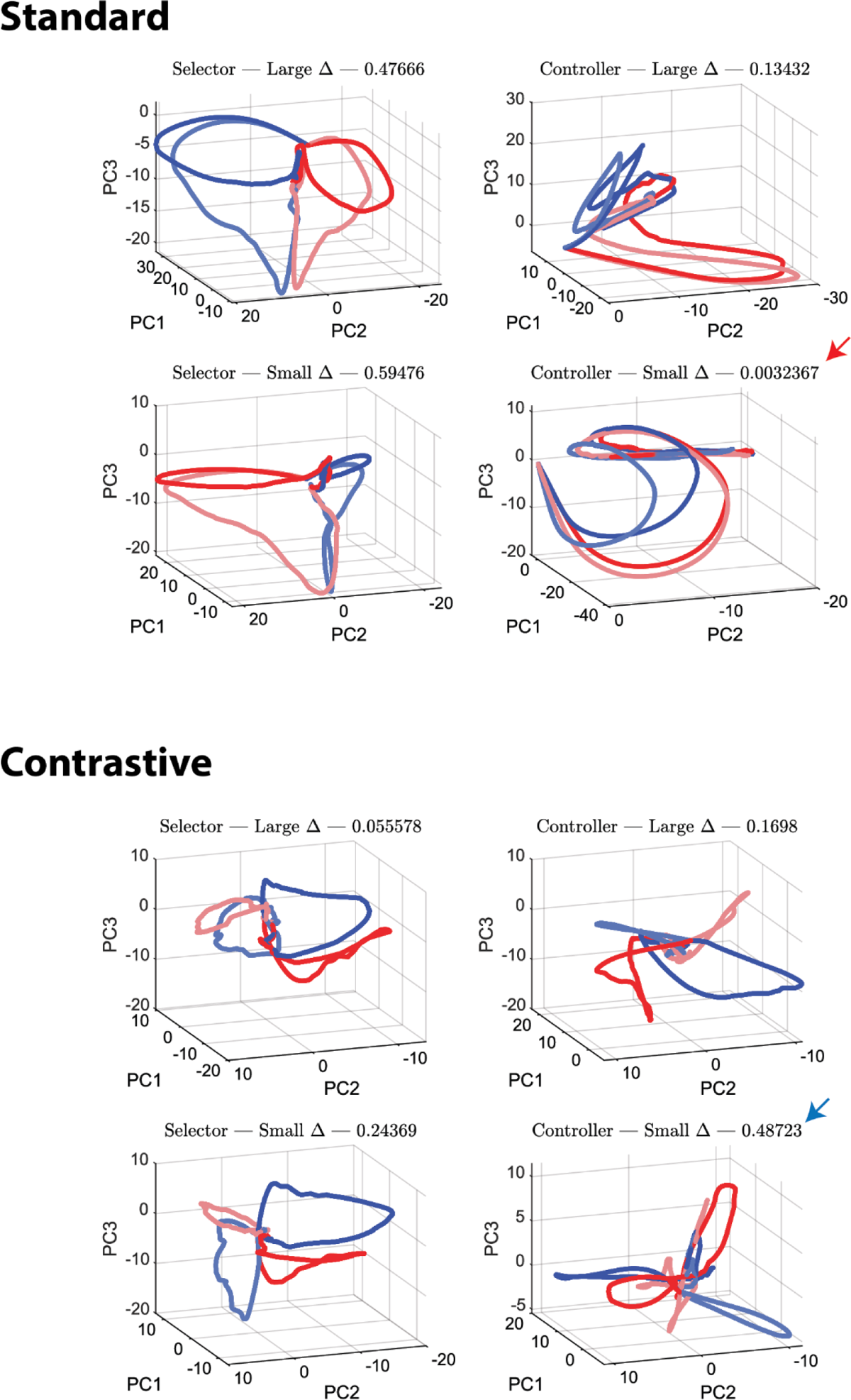
Differential activity patterns that can be revealed by cPCA The encoding model from Supp Fig. 1 can be used to provide ground truth for the conjecture that contrastive PCA can reveal separation in population dynamics as a function of action, even from small actions that differ little in the underlying parameters/encoding. Upper panels show standard PCA projection for example model run. Lower panels show contrastive PCA projection for analogous model runs. Arrow heads indicate the large change in rotation specifically for the Controller Small Delta simulations. Full quantification of distribution of rotation values shown in main text and Figure 2.

**Supp. Fig. 4.**
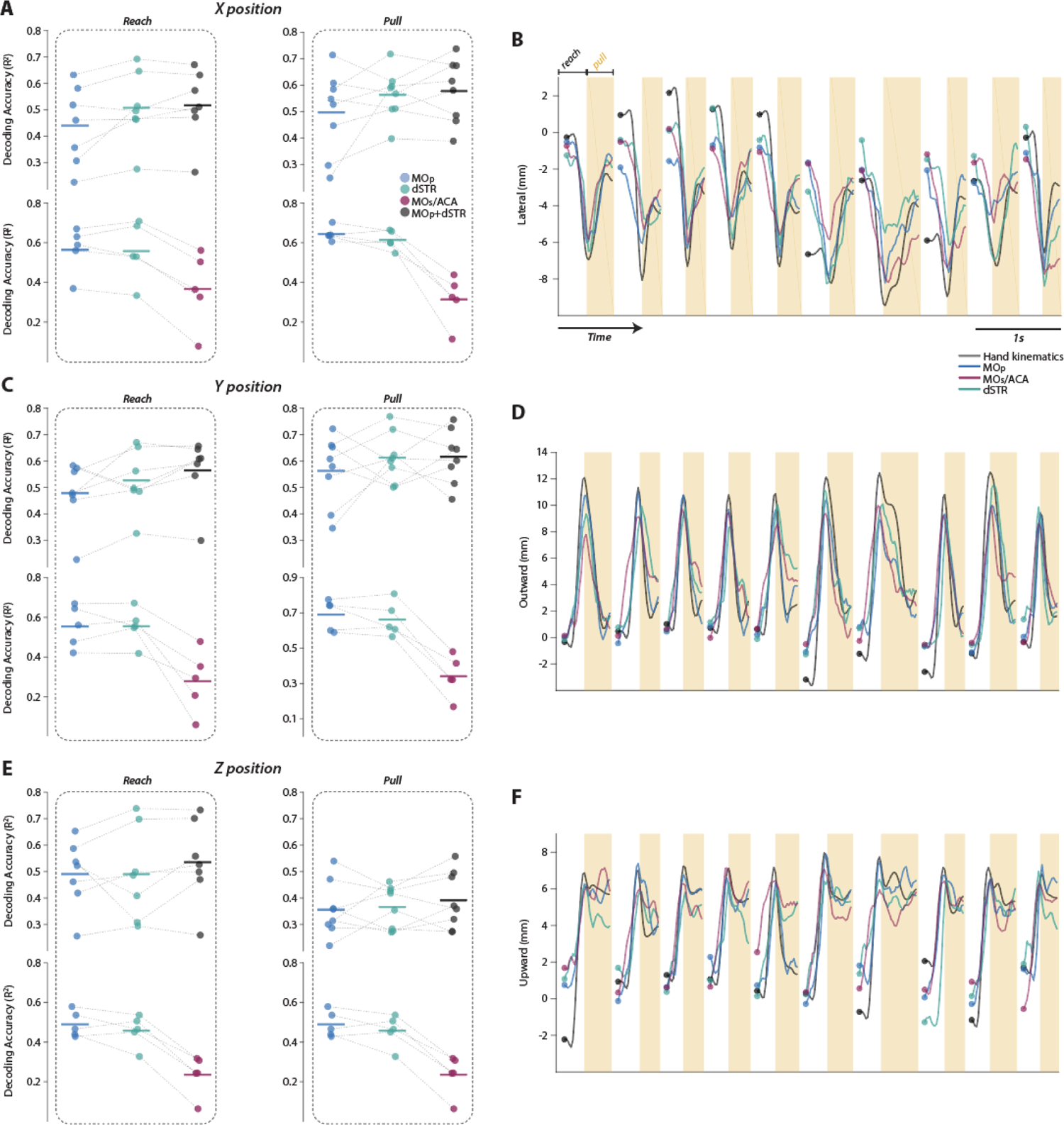
Continuous decoding of 3D hand trajectories The Kalman filter was used to decode 3D hand trajectories based on the neural peri-movement responses with cross-validation and resampling to match the number of neurons across regions. **A** Accuracies (R^2^) of decoding hand trajectories on the x axis (left-right) were compared across regions during reach and pull phases. **B** Ten representative hand trajectories on the x axis decoded from each region were superimposed with the observed hand trajectories (black). C Formatted same as A but accuracies are of trajectories on the y axis (forward-backward). **D** Formatted the same as B but trajectories represent hand positions on the y axis. **E** Formatted same as A but accuracies are of trajectories on the z axis (upward-downward). F Formatted the same as B but trajectories represent hand positions on the z axis.

**Supp. Fig. 5.**
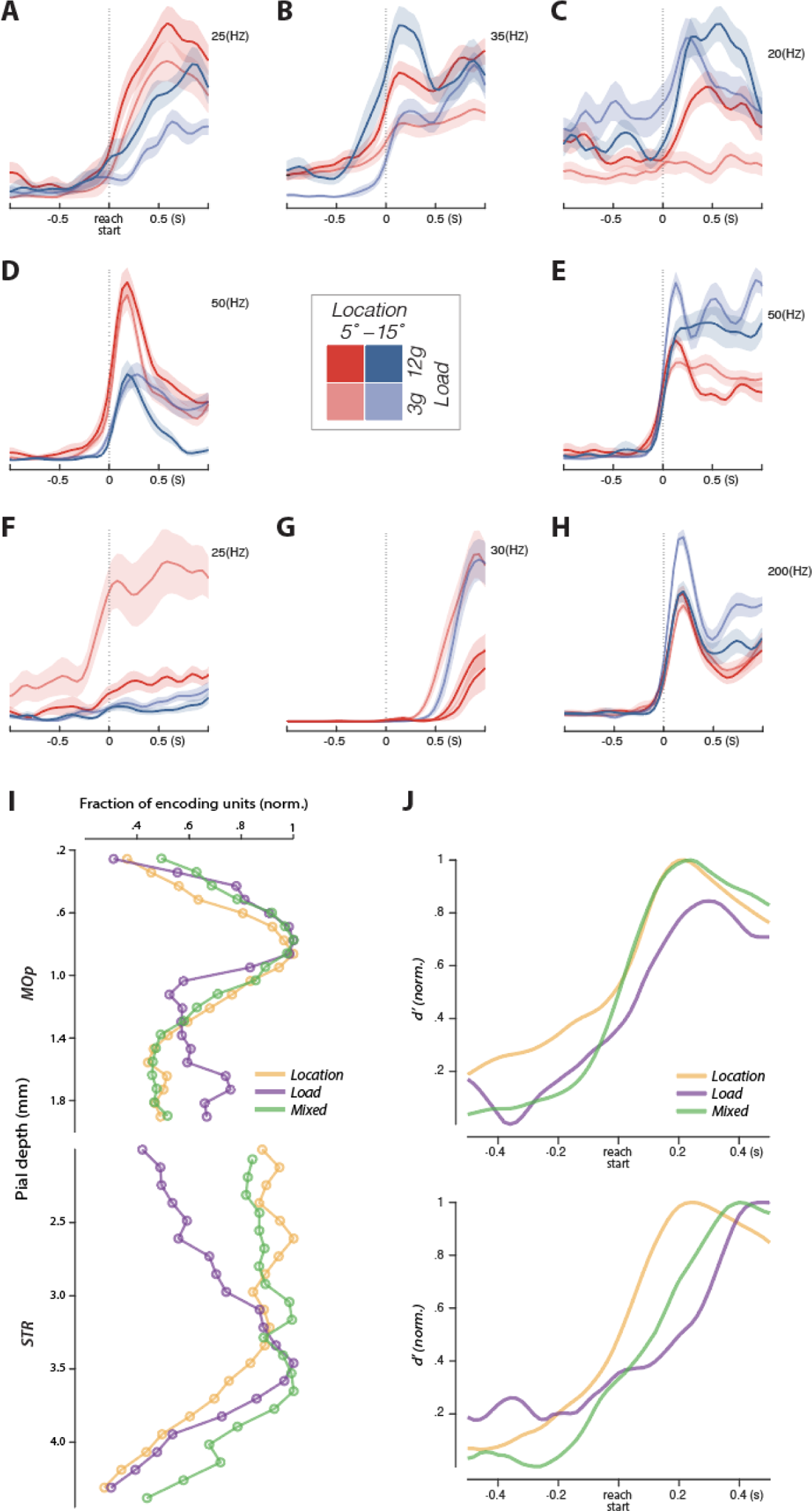
More details on MOp and dSTR single neuron encoding of trial conditions **A-H** Individual neurons were characterized by their responses to joystick location and load combinations. **A** An example neuron with preferred response to both leftward location and high joystick load. **B** An example neuron with preferred response to high load with little sensitivity to target location. **C** An example neuron with preferred response to both rightward location and high load. **D** An example neuron with preferred response to leftward target location with little sensitivity to load. **E** An example neuron with preferred response to rightward target location with little sensitivity to load. **F** An example neuron with preferred response to both leftward location and high load. **G** An example neuron with preferred response to low load with little sensitivity to target location. **H** An example neuron with preferred response to both rightward location and low load. **I-J** Some individual neurons purely encoded joystick ‘location’ (D & E) or ‘load’ (B & G), whereas other neurons displayed ‘mixed’ encoding of both (A, C, F, H). The spatiotemporal distribution of the three encoding types were examined. **I** Spatial distribution of ‘location’, ‘load’, and ‘mixed’ encoding neurons along the corticostriatal depth. Fractions of encoding neurons were normalized to the maximum fraction of each encoding group. **J** Temporal responses properties of the three encoding groups. The *d’* scores were normalized to the maximum score of each encoding group.

**Supp. Fig. 6.**
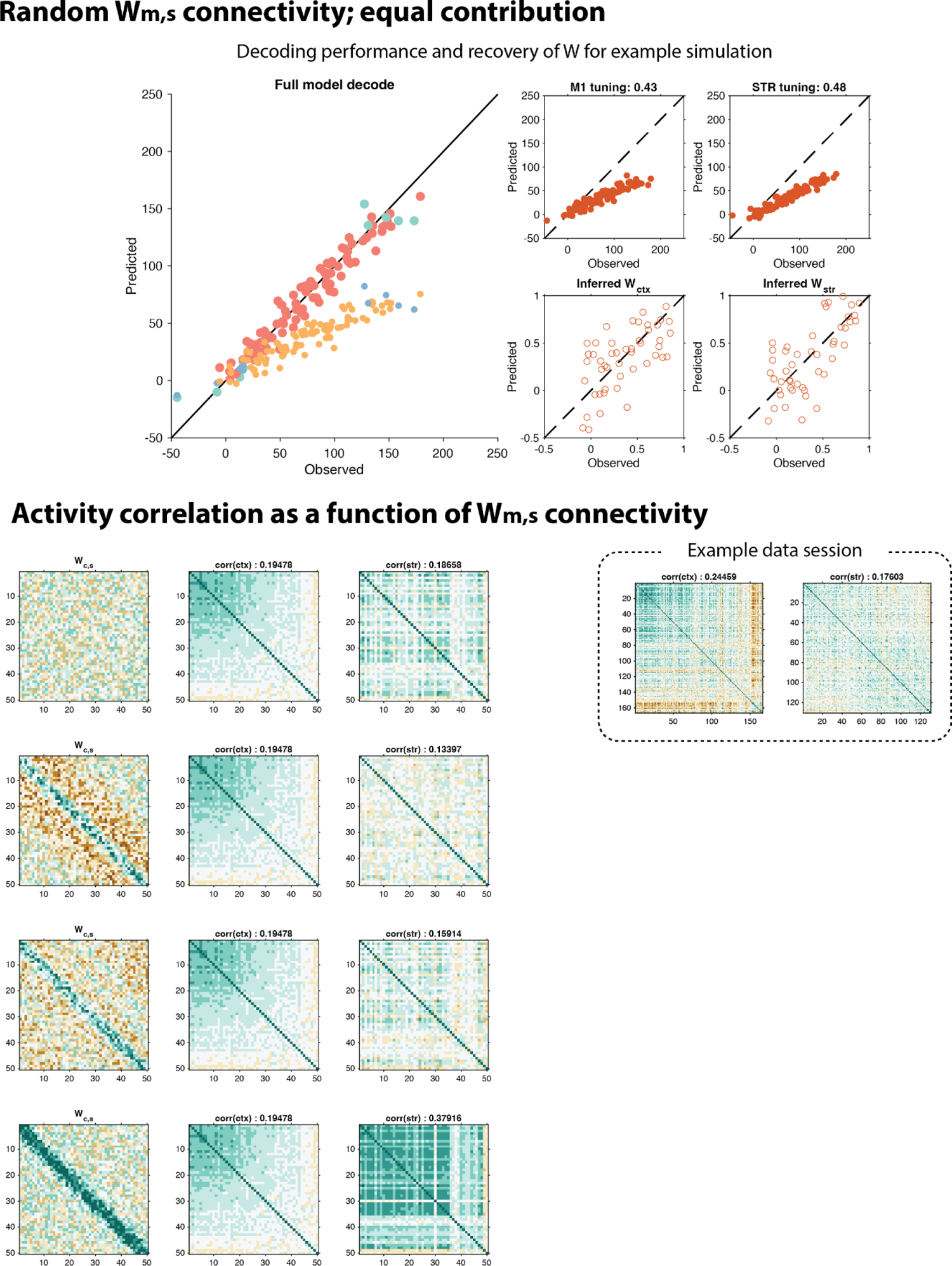
Elaboration on the 2 layer encoding model details The 2 layer encoding model described in the main text is sufficient to recover equivalent decoding contributions from motor cortex (‘m’ layer) and striatum (‘s’ layer) under the assumption that the two are independent (upper panels) or coupled by a random Wm,s matrix. To illustrate how s correlations vary as a function of Wm,s we illustrate random (top), center_surround (second), distributed center surround (third, used for calculations in main text), and topographic excitatory (bottom). An example data session pairwise correlations of integrated response during pull phase correlations are shown at right (distribution shown in main text).

---

**Supp. Video 1.** Example reach-to-pull trials https://drive.google.com/open?id=1ZL7tRgWUHBNzKRpExP8gIzYQgXmWGGhc&usp=drive_fs

**Left:** Reproduction of 10 consecutive reach-to-pull trials (joystick location: 5°, load: 3g) with 0.5x playback speed. Right: Reproduction of hand velocities decoded from simultaneously recorded MOp (blue) and STR neural populations (green) are superimposed with the actual hand velocity inferred from DeepLabCut (black).

## Footnotes

1 Quantitatively, the active ensembles must be less correlated than the ensembles in the input as in (*28*). In many accounts, including (*28*), the representations proposed are only static patterns associated with an action (e.g. (*13*, *28*)); however, this is clearly inconsistent with the complex temporal dynamics of dSTR (and MOp) activity during movement (*32*, *34*, *39*). Thus, to ‘steelman’ the selection model we entertain a stronger version including temporal dynamics (*6*).

2 Dimensions of activity other than leading PCs provide a robust encoding of continuous parameters; later we consider how to find such activity dimensions in the case of a specification encoding model.

## Notes

https://dudlab.notion.site/Conjoint-specification-of-action-in-neocortex-and-striatum-cebef9fc306947139f0ee70ce2dadeaa

### Competing Interest Statement

The authors have declared no competing interest.

